# Impact of pH and temperature in dairy processing on the infectivity of H5N1 avian influenza viruses

**DOI:** 10.1101/2025.03.21.644501

**Authors:** Nicole-Lenz-Ajuh, Leonie Rau, Lisa Butticaz, Bettina Zimmer, Vincent Beuret, Florian Loosli, Jan-Erik Ingenhoff, Barbara Wieland, Gert Zimmer

**Affiliations:** Agroscope, Schwarzenburgstrasse 161, CH-3097 Liebefeld, Switzerland; Institute of Virology and Immunology IVI, Sensemattstrasse 293, CH-3147 Mittelhäusern, Switzerland; Department of Infectious Diseases and Pathobiology, Vetsuisse Faculty, University of Bern, Länggassstrasse 120, CH-3012 Bern, Switzerland

**Keywords:** Raw milk, lactic acid fermentation, soft cheese, semi-hard cheese, yoghurt, pasteurization, hemagglutinin

## Abstract

Highly pathogenic avian influenza viruses (HPAIV) of subtype H5N1 (clade 2.3.4.4b) have crossed the species barrier and caused a mastitis-like infection in dairy cows. The high levels of infectious virus found in the milk raised considerable concerns about the safety of raw milk products. This study examined the effect of temperature and pH on the stability of HPAIV and low-pathogenic avian influenza viruses (LPAIV). We found that H5N1 HPAIV remained infectious in milk at 4°C for four weeks, with slow decreases at 21°C, and complete inactivation at 37°C after four weeks. H5N1 LPAIV was stable at 50°C for 30 minutes but inactivated at higher temperatures (55°C for 10 minutes, 60°C for 1 minute, or 72°C for 30 seconds). At pH levels between 6 and 10, the virus remained stable but was partially inactivated at pH 5.0 and completely inactivated at pH 4.0. During yogurt production, H5N1 LPAIV was completely inactivated when the pH reached 4.3. In cheese production, the lowest pH reached was between 5.0 and 5.3. When H5N1 LPAIV was incubated with soft and semi-hard cheese for one day at 4 °C, infectious virus titers decreased by 5.1 and 3.9 log_10_, respectively. When H5N1 LPAIV was incubated with buffer adjusted to pH 5.0, infectious virus titer dropped by only 3.3 log_10_, suggesting that, alongside pH, other processes of cheese ripening likely influence virus stability. In conclusion, H5N1 avian influenza viruses are largely inactivated during lactic acid fermentation of raw milk. Future studies will assess the required cheese ripening time for complete inactivation.

## 1. Introduction

In February 2024, a milk drop syndrome was recorded in dairy cows in Texas, USA. In March 2024, highly pathogenic avian influenza virus (HPAIV) of subtype H5N1 clade 2.3.4.4b, genotype B3.13 was identified as the etiological agent of the syndrome (Mostafa et al., 2024; Webby and Uyeki, 2024). Although this avian influenza virus did not show mutations characteristic of adaptation to a mammalian host (Hu et al., 2024), it was still designated “bovine H5N1” (Eisfeld et al., 2024). The infected animals suffered from a mastitis-like disease with fever, low appetite, reduced milk production, and abnormal milk consistence (Halwe et al., 2025). Animals generally recovered only very slowly from the disease and milk yield remained low for at least 4 weeks (Caserta et al., 2024).

The virus was found to replicate predominantly in the mammary glands of lactating cows with high infectious virus titers shed into the milk (≥10^8^ ffu/ml) (Halwe et al., 2025). The virus was further transmitted between animals by contaminated milking equipment (Le Sage et al., 2024), the role of other transmission routes remains unclear. Moreover, virus spill over to cats, mice, and poultry may have occurred (Mainenti et al., 2025; Mostafa et al., 2024; Naraharisetti et al., 2025). In addition, several farm workers who had been in contact with infected cows got infected. Fortunately, most of these persons displayed only mild symptoms such as conjunctivitis (Garg et al., 2024; Uyeki et al., 2024). A serological survey among 150 bovine veterinary practitioners who have been exposed to infected cows revealed that about 2% of them seroconverted to H5N1 (Leonard et al., 2025).

Within a year, bovine H5N1 affected approximately 983 herds in 17 US states by March 2025 (USDA, 2025). Recently, genotype D1.1 of H5N1 HPAIV (clade 2.3.4.4b) had also been detected in dairy cattle in the United States, indicating that H5N1 HPAIV have been transmitted from wild birds to dairy cows at least two times (Pekar et al., 2025). There is also experimental evidence that European H5N1 HPAIV (clade 2.3.4.4b) isolates can cause a similar disease in lactating cows suggesting that the phenomenon is not unique to the American genotypes B3.13 and D1.1, but might be rather a more general feature of H5N1 HPAIV (clade 2.3.4.4b) (Halwe et al., 2025).

Since infected dairy cows were found to shed large amounts of infectious H5N1 HPAIV into the milk, it was not surprising that H5N1 genomic RNA was detected in around 17.4% of retail dairy products in the USA (Suarez et al., 2025). However, infectious virus was not detected in pasteurized milk (Spackman, Jones, et al., 2024), and recent studies confirmed that H5N1 HPAIV is inactivated by treatment for 15 seconds at 72°C (pasteurization) (Guan et al., 2024). Feeding of virus-contaminated unpasteurized milk to mice resulted in lethal infection of mice and cats (Burrough et al., 2024; Eisfeld et al., 2024). A recent study in macaques showed that non-human primates reacted differentially, as they did not show any severe symptoms upon oral infection with bovine H5N1 (Rosenke et al., 2025). Nevertheless, there are still considerable concerns about the safety of milk products since different types of cheese and other dairy products are produced from raw milk.

Since lactic fermentation is a key process in cheese production, the aim of the present study was to evaluate the impact of two important parameters in cheese production, pH and temperature, on infectivity of H5N1 avian influenza viruses. As storage of contaminated milk at certain temperatures can have an impact on virus infectivity as demonstrated for Zika virus, Rift valley fever virus, hepatitis C virus, and tick-borne encephalitis virus (Dawes et al., 2025; Offerdahl et al., 2016; Pfaender et al., 2013; Pfaender et al., 2017), this parameter was investigated in the present study as well. Our findings may provide valuable information that can help with the risk assessment of raw milk products.

## 2. Materials and methods

### 2.1. Cells

Madin-Darby canine kidney (MDCK) type II cells were kindly provided by Georg Herrler (University of Veterinary Medicine, Hannover, Germany) and maintained with minimum essential medium (MEM, Life Technologies, Zug, Switzerland) supplemented with 5% of fetal bovine serum (FBS; Pan Biotech, Aidenbach, Germany).

### 2.2. Viruses

HPAIV A/Cattle/Texas/063224-24-1/2024 (H5N1), genotype B3.13 (GISAID accession number: EPI_ISL_19155861) was kindly provided by Diego Diel (Cornell University, Ithaca, NY, USA). This virus was originally isolated from the milk of infected dairy cows in Texas, USA (Caserta et al., 2024). A virus stock was produced on MDCK cells and stored in aliquots at −70 °C. Infectious virus titers were determined by endpoint titration on MDCK cells and expressed as tissue infectious dose 50% (TCID_50_).

The LPAIVs A/duck/Hokkaido/Vac-1/2004 (H5N1) (Soda et al., 2008) and A/duck/Potsdam/1402/1986 (H5N2) (GenBank accession numbers: CY00577 – CY005783, CY014642) were kindly provided by Yoshiro Sakoda (Hokkaido University, Sapporo, Japan) and Timm Harder (Friedrich-Loeffler-Institute, Greifswald - Insel Riems, Germany), respectively. Both viruses were propagated for 2 days at 37 °C in the allantoic cavities of 10-day-old embryonated specific pathogen-free (SPF) chicken eggs.

The chimeric vesicular stomatitis virus VSVΔG(HA:NA:GFP) encoding the HA and NA glycoproteins of HPAIV A/Pelican/Bern/1/2022 (H5N1), clade 2.3.4.4b (GISAID accession numbers accession numbers EPI3526757 and EPI3526758), along with green fluorescent protein (GFP) has been generated according to a previously reported method (Thompson et al., 2023). The virus was propagated on MDCK cells and aliquots stored at −70 °C.

### 2.3. Virus titration

Infectious virus titers of HPAIV were determined on MDCK cells by limiting dilution. To this end, MDCK cell monolayers in 96-well tissue culture plates were incubated in quadruplicates with serially diluted virus (100 μL/well) for 2 days at 37 °C. The cells were washed once with PBS (200 μL/well) and fixed for 1 hour at 21 °C with 4 % of buffered formalin containing 0.1 % (w/v) crystal violet. The plates were washed with tap water and dried. Virus titer was calculated using the Spearman-Kärber method and expressed as tissue infectious dose 50 % (TCID_50_) (Ramakrishnan, 2016). The titer of the virus stock used in the present study was 10^8^ TCID_50_/mL.

For determination of infectious LPAIV titers MDCK cells grown in 96-well tissue culture plates were inoculated in duplicate with 40 µL per well of serial 10-fold virus dilutions for 1 hour at 37 °C. Thereafter, 160 µL of MEM containing 1 % methylcellulose was added to each well, and the cells incubated for 24 hours at 37 °C. The cells were fixed for 30 minutes with 4% formalin in PBS, permeabilized with 0.25 % (v/v) of Triton X-100, and incubated for 60 minutes with a monoclonal antibody directed to the influenza nucleoprotein (American Type Culture Collection, Manassas, Virginia, USA, ATCC HB-65, clone H16-L10-4R5), diluted 1:50 with PBS, and subsequently for 60 minutes with goat anti-mouse IgG conjugated with Alexa Fluor-488 (diluted 1:500 in PBS; Life Technologies, Zug, Switzerland). Infected cells were detected by fluorescence microscopy (Observer Z1 microscope, Zeiss, Feldbach, Switzerland), and infectious virus titers were calculated and expressed as focus-forming units per milliliter (FFU/mL). The titer of the virus stocks used in the present study were 5x 10^8^ FFU/mL for both LPAIV H5N1 and H5N2. VSVΔG(HA:NA:GFP) was titrated in an analogous manner using the GFP reporter for detection and enumeration of infected cells.

### 2.4. Thermal stability of H5N1 avian influenza viruses

Fresh raw milk was obtained from a regional dairy farm and heated for 10 minutes at 90 °C to prevent bacterial growth during long-term incubation experiments. The heat-inactivated milk (450 µL) was spiked with 50 µL of either HPAIV A/Cattle/Texas/063224-24-1/2024 (H5N1) or LPAIV A/duck/Hokkaido/Vac-1/2005 (H5N1) and incubated for up to 28 days at either 4 °C, 21 °C, or 37 °C. Every 7 days, three samples per temperature of incubation were frozen at −70 °C. Finally, infectious titers were determined by virus titration on MDCK cells (see section 2.2.).

For treatment of LPAIV H5N1 virus at temperatures ≥ 37 °C, 80 µL of either heat-inactivated milk or 80 µL of MEM medium were spiked with 20 µL of A/duck/Hokkaido/Vac-1/2005 (H5N1) and incubated in thin-walled 0.2 ml PCR tubes (Eppendorf SE, Hamburg, Germany) for the indicated times and temperatures using a T100 Thermal Cycler (BioRad, Cressier, Switzerland) with the lid heated to 105 °C. Infectious titers were determined by virus titration on MDCK cells (see section 2.2.).

### 2.5. Polykaryon formation assay

MDCK cells were grown on cover slips (12-mm in diameter) and infected with VSVΔG(H5;N1:GFP) using a multiplicity of infection (MOI) of 0.01 FFU/cell. At 20 hours post infection, the cells were shortly rinsed with MES buffer (50 mM 4-morpholine ethane sulfonic acid, 150 mM NaCl, 1 mM CaCl_2_, 1 mM MgCl_2_), adjusted to pH 5.0 – 6.2, and incubated in the respective buffer for 10 minutes at 37 °C. Thereafter, the cells were washed once with MEM medium and incubated for 2 hours at 37 °C in MEM medium supplemented with 5 % FBS. The cells were washed once with PBS and fixed for 30 minutes at 21 °C with PBS containing 4 % formalin. For staining of nuclei, the cells were incubated for 5 minutes at 37°C with 0.1 µg/mL of 4′,6-diamidino-2-phenylindole (DAPI, Merck KGaA, Darmstadt, Germany) and mounted in ProLong Gold Antifade Mountant (Thermo Fisher). Image acquisition was performed using the LCI Plan-NEOFLUAR 63×/1.3 water immersion objective of an Observer Z1 fluorescence microscope (Zeiss).

### 2.6. Analysis of virus stability at different pH values

For the treatment of viruses at different pH values, the following buffer systems were used: 50 mM Na_2_HPO4/citric acid, 150 mM NaCl, pH 2.0 - 5.0; 50 mM MES, 150 mM NaCl, pH 5.0 – 7.0; 50 mM Tris/HCl, 150 mM NaCl, pH 7.0 - 10.0. To 450 μL of the respective buffer 50 μL of either HPAIV A/Cattle/Texas/063224-24-1/2024 (H5N1), LPAIV A/duck/Hokkaido/Vac-1/2005 (H5N1) or LPAIV A/duck/Potsdam/1402/1986 (H5N2) were added and incubated for 30 minutes at 21 °C. To replace the buffer with cell culture medium, each incubation (500 μL) was loaded on a Sephadex G-25 columns (PD MiniTrap G-25, Cytiva Europe, Freiburg, Germany) that has been equilibrated with MEM medium containing 5% FBS and peniciliin/streptomycin (Life Technologies, Zug, Switzerland). The virus was eluted from the columns by adding 1 mL of MEM medium containing 5% of FBS and penicillin/streptomycin and subsequently titrated on MDCK cells (see section 2.2.).

### 2.7. Production of yoghurt using LPAIV H5N1-spiked raw milk

For production of yoghurt, the Jog BL1 starter culture (Liebefeld Kulturen AG, Switzerland), consisting of a mixture of *Streptococcus thermophilus*, *Lactobacillus delbrueckii* ssp. *bulgaricus* and *Lactobacillus delbrueckii* ssp. *lactis*, was prepared according to the instructions of the manufacturer. Briefly, 1.2 % of the BL1 culture was incubated for 3.5 hours at 42 °C in ultra-heated skimmed milk (Cremo, Switzerland) and then stored at 4 °C until further use. Yoghurt was produced using 5 mL aliquots of fresh raw milk with 3 % skimmed milk powder (Rapilait, Migros) with or without 1 % of the prepared BL1 culture and 0.5 ml of LPAIV A/duck/Hokkaido/Vac-1/2005 (H5N1). The virus-spiked yoghurt was incubated in a water bath at 42 °C until the pH was in the range of 4.0 - 4.6 (after ca. 5 hours of incubation). The fermentation process was terminated by incubating the yoghurt overnight at 4 °C. The yoghurt (5 mL) was mixed with 5 mL of MEM medium and centrifuged (3000 x g) for 15 minutes at 4 °C to pellet precipitated protein. Finally, the supernatant was passed through a 0.2 μm pore size filter and stored at −70° prior to virus titration (see section 2.2.) or RT-qPCR analysis (see section 2.9.).

### 2.8. Incubation of LPAIV H5N1 with freshly manufactured raw milk cheese

For soft mini-cheese production, the BAMOS starter culture (Liebefeld Kulturen AG, Switzerland), consisting of a mixture of *Streptococcus thermophilus* and *Lactobacillus delbrueckii* ssp. *bulgaricus,* was prepared according to the instructions of the manufacturer. Briefly, 0.6 % of the BAMOS culture was incubated for 4 hours at 40 °C in ultra-heated skimmed milk (Cremo, Switzerland) and stored at 4 °C until further use. Soft cheese was produced by incubating 30 mL of fresh raw milk with 0.5 % of starter culture for 30 minutes at 35 °C. Thereafter, 0.025 % of beef rennet (Winkler GR orange, Switzerland) was added and incubated in a water bath at 35 °C for 30 minutes. The curd was cut into pieces of roughly 15 mm^3^ and incubated at 35 °C for 40 minutes with 6 mL of tap water. Subsequently, the milk was centrifuged (3000x g) for 20 minutes at 21 °C, the supernatant (whey) aspirated, and the precipitated material incubated at 35 °C until the pH was in the range pH 5.0 – pH 5.3. Finally, the soft cheese was refrigerated at 4°C for 7 days.

For semi-hard mini-cheese production, the RMK 302 starter culture (Liebefeld Kulturen AG, Switzerland), consisting of a mixture of *Lactobacillus delbrueckii* ssp. *lactis* and *Streptococcus thermophilus* was prepared according to manufacturer’s instructions. Briefly, 0.2 % of the RMK 302 starter culture was incubated in ultra-heated skimmed milk (Cremo, Switzerland) for 6 to18 hours at 38 °C and stored at 4 °C until further use. Semi-hard cheese was produced using 30 mL aliquots of fresh raw milk mixed with 0.5 % of the starter culture and incubated for 30 minutes at 35 °C. Thereafter, 0.025 % of beef rennet (Winkler GR orange, Switzerland) was added and incubated for 30 minutes at 35 °C. The curds were cut to pieces of 4 - 8 mm^3^ to which 4.5 mL of tap water were added. The temperature was increased from 35 °C to 45 °C within 15 minutes and the temperature was held at 45 °C for another 15 minutes. Precipitated proteins were pelleted by centrifugation (3000x g, 20 minutes) and incubated at 40 °C until the pH dropped to 5.0 - 5.3. The semi-hard cheese was stored refrigerated at 4°C.

For incubation of LPAIV A/duck/Hokkaido/Vac-1/2005 (H5N1) with 7days-old soft cheese or semi-hard cheese, about 0.6 g of the respective cheese was cut into small pieces of 0.5 mm^3^ and suspended with 0.9 mL of sterile water. Measurement of the proton concentration showed that the pH was in the range of 5.0 – 5.3. To the cheese 0.1 mL of virus stock was added and incubated for 24 hours at 4°C. Subsequently, 0.5 mL of the incubated virus suspension was loaded onto Sephadex G-25 columns and eluted in 1 mL of MEM cell culture medium supplemented with 5% FBS and penicillin/streptomycin.

### 2.9. Detection of viral RNA by RT-qPCR

RNA was extracted from 200 μL of spiked raw milk using the NucleoMag Vet kit (Macherey-Nagel AG, Oensingen, Switzerland according to the manufacturer’s protocol. Reverse transcription from RNA to cDNA and real-time quantitative PCR (qPCR) were performed on the QuantStudio 5 real-time PCR system (Thermo Fisher Scientific) using the AgPath-ID One-Step RT-PCR kit (Life Technologies, Zug, Switzerland) and segment 7-specific oligonucleotide primers and probe (Hofmann et al., 2008; Spackman et al., 2002). Data were acquired and analyzed using the Design and Analysis Software v1.5.2 (Thermo Fisher Scientific).

### 2.10. Statistical analysis

Statistical analysis was performed using GraphPad Prism 10, version 10.1.2 (GraphPad Software, Boston, Massachusetts, USA). Unless noted otherwise, the results are expressed as mean ± standard deviation (SD). Specific statistical tests such as the one-way or two-way ANOVA test were used to assess significant differences in serum antibody responses in vaccinated animals as indicated in the figure legends. P values < 0.05 were considered as significant.

## 3. Results

### 3.1. Impact of long-term storage of HPAIV and LPAIV H5N1 in bovine milk

First, we investigated the effect of long-term storage of bovine milk spiked with H5N1 avian influenza viruses on infectious virus titers. We incubated either HPAIV A/Cattle/Texas/063224-24-1/2024 (H5N1) or LPAIV A/duck/Hokkaido/Vac-1/2004 (H5N1) with bovine milk for four weeks at either 4 °C, 21 °C or 37 °C. Aliquots of the virus-spiked milk were taken at weekly intervals and the infectious virus titers on MDCK cells were determined by endpoint titration (**Fig. 1**). Both H5N1 HPAIV and H5N1 LPAIV were stable when incubated for four weeks at 4 °C and showed no significant loss of infectivity during this time. However, when the viruses were incubated with milk at 21 °C, their infectivity decreased over time. While H5N1 LPAIV had a tenfold reduced infectious titer after four weeks, the infectious titer of H5N1 HPAIV was reduced by about 2.5 log _10_ at the end. Incubation of the viruses at 37 °C had a stronger effect on virus infectivity. While the infectious titer of H5N1 HPAIV fell below the detection limit after three weeks of incubation, infectious H5N1 LPAIV was still detected after four weeks of incubation, albeit at significantly reduced levels. These findings suggest that there may be strain-specific differences in virus stability. However, at 4 °C, the infectious titers of both virus strains remained high, with no signs of inactivation by milk components.

**Fig. 1.**
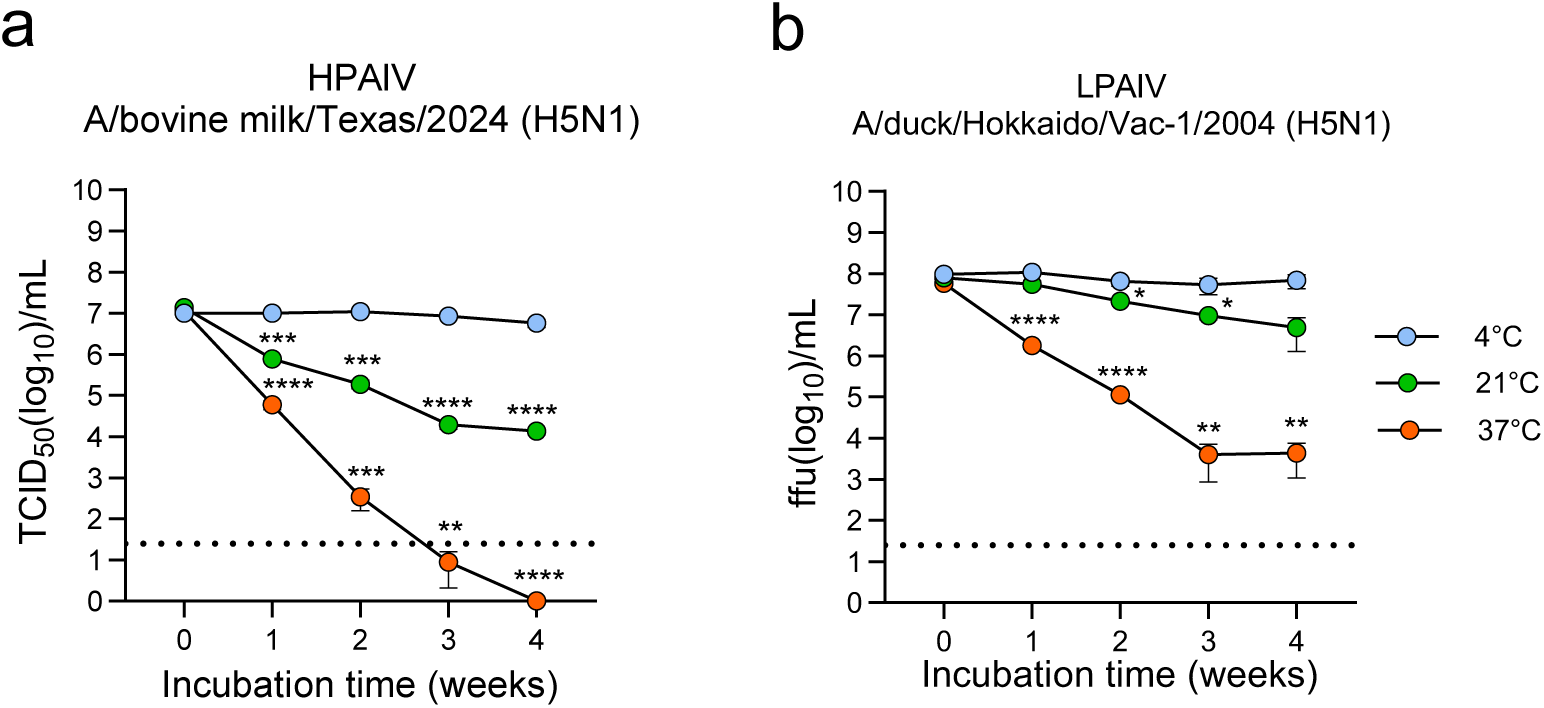
Impact of long-term storage of H5N1 in bovine milk. Raw milk was heated for 10 minutes at 95 °C, spiked with either H5N1 HPAIV (**a**) or H5N1 LPAIV (**b**), and incubated for up to 4 weeks at either 4 °C, 21 °C, or 37 °C. At the indicated time points, the samples were frozen at −70 °C. After the last sample was collected, all samples were thawed and infectious virus titers determined on MDCK cells by end point dilution. Mean titers and standard deviations of three incubation experiments are shown. The two-way ANOVA with Tukey’s multiple comparison test was used to assess significantly different infectious titers (\**p* < 0.05, \*\**p* < 0.01, \*\*\**p* < 0.001, \*\*\*\**p* < 0.0001).

### 3.2. Thermal stability of LPAIV H5N1 at elevated temperatures

To investigate viral stability at higher temperatures, H5N1 LPAIV was diluted in MEM cell culture medium or bovine milk and incubated for 30 minutes at 4 °C, 37 °C, 40 °C, 45 °C, 50 °C, 55 °C or 60 °C. Subsequently, the infectious viral titer on MDCK cells was determined by endpoint titration (**Fig. 2a**). We found that H5N1 LPAIV was as stable in bovine milk at 37 °C as it was at 4 °C, while at 40 °C and 45 °C there was a small reduction in infectious virus titer (2-fold and 3.5-fold, respectively). On the other hand, a more pronounced reduction in viral titer was found when the virus was incubated at 50 °C (35 - fold reduction). Of note, no infectious virus was detectable at all when the virus was incubated for 30 minutes at 55 °C or 60 °C. A very similar inactivation pattern was found for H5N1 LPAIV incubated with MEM medium (**Fig. 2a**). However, the infectious virus titers remained at generally higher levels compared to virus incubated with milk, suggesting that H5N1 LPAIV is less stable in bovine milk. To determine the minimum incubation period leading to complete virus inactivation, we incubated H5N1 LPAIV with bovine milk at 55°C or 60°C for shorter periods. We observed that infectious virus titers fell below the detection limit when the virus was treated for 10 minutes at 55 °C or for 1 minute at 60 °C (**Fig. 2b, c**). Likewise, the virus was completely inactivated when incubated for 30 seconds at 72 °C (**Fig. 2d**). When the virus was incubated with bovine milk at 72 °C for 15 seconds, a titer reduction by 4.5 log_10_ was achieved, which is consistent with previous reports showing that pasteurization can efficiently inactivate bovine H5N1 HPAIV in milk (Alkie et al., 2025; Cui et al., 2024; Spackman, Anderson, et al., 2024).

**Fig. 2.**
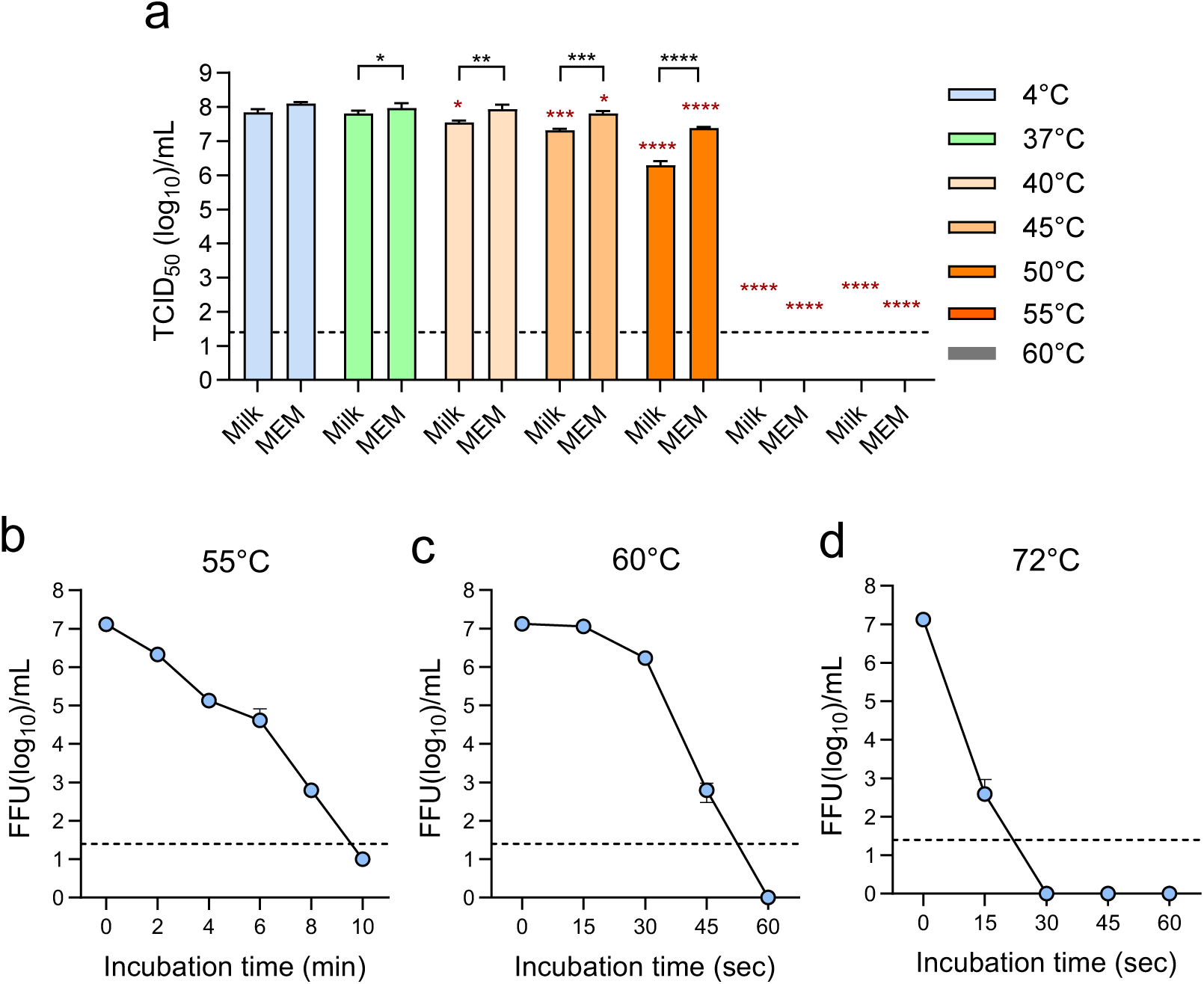
Thermal stability of LPAIV H5N1. (**a**) Bovine milk or MEM medium were spiked with LPAIV A/duck/Hokkaido/Vac-1/04 (H5N1) and incubated for 30 minutes at the indicated temperatures. (**b-d**) Bovine milk was spiked with A/duck/Hokkaido/Vac-1/04 (H5N1) and incubated for the indicated times at either 55 °C (**b**), 60 °C (**c**) or 72 °C (**d**). Infectious virus titers were determined on MDCK cells by end point titration. The interrupted line indicates the limit of detection of infectious virus. Mean values and standard deviations of three inactivation experiments are shown. Significantly differences in infectious virus titers were calculated using the two-way ANOVA test. Significant differences between spiked milk and MEM are indicated by black asterisks (Sidak’s multiple comparisons) and significant differences between elevated temperatures and 4°C are indicated by red asterisks (Tukey’s multiple comparisons); \**p* < 0.05; \*\**p* < 0.01, \*\*\**p* < 0.001, \*\*\*\**p* < 0.0001.

### 3.3. The HA of H5N1 HPAIV clade 2.3.4.4b initiates membrane fusion at mildly acidic conditions

Influenza A viruses enter cells by receptor-mediated endocytosis and fusion of the viral envelope with the endosomal membrane. Membrane fusion is mediated by hemagglutinin (HA), the major viral envelope glycoprotein, which is usually in a metastable conformation. The acidification of the endosomal lumen triggers a conformational change in HA which initiates the membrane fusion process. As this conformational change is irreversible, a premature exposure of influenza A viruses to low pH can result in HA denaturation and complete loss of infectivity. To investigate at which pH threshold value, the HA of H5N1 HPAIV would start triggering membrane fusion, we performed a polykaryon formation assay. To this end, we infected MDCK cells with a recombinant vesicular stomatitis virus vector encoding the HA and NA antigens of H5N1 HPAIV (clade 2.3.4.4b) along with the green fluorescent protein (GFP) reporter. At 16 hours post infection, the cells were shortly incubated with buffers adjusted to different pH values and subsequently incubated for 2 hours at 37°C with cell culture medium. The cells were fixed and analyzed for the formation of polykaryons (syncytia) (**Fig. 3a**). The results indicate that mildly acidic conditions (pH of 6.0 - 6.1) are sufficient to trigger a conformational change in the HA of H5N1 HPAIV (clade 2.3.4.4b), which leads to cell-to-cell fusion.

**Fig. 3.**
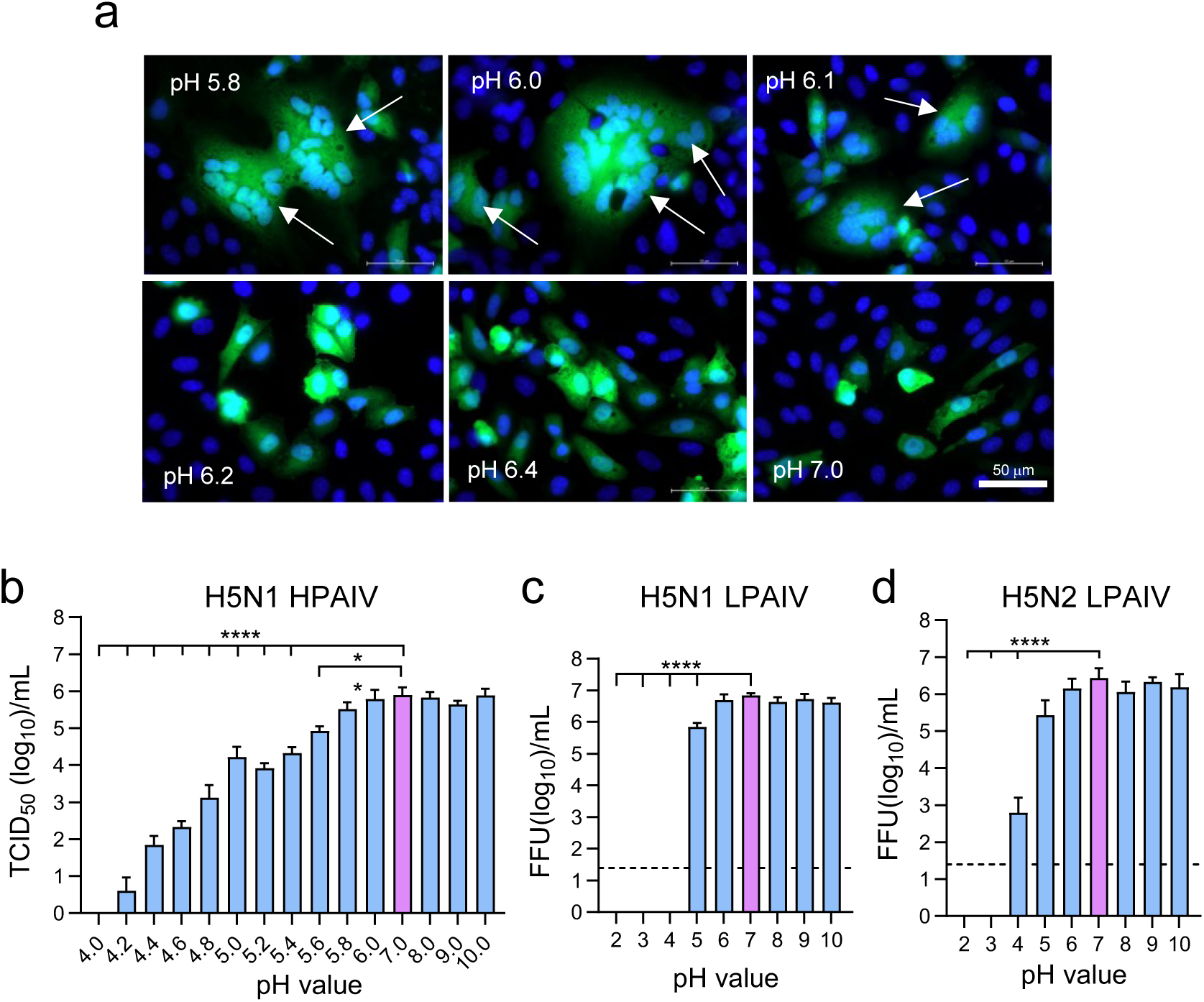
Acid stability of H5 avian influenza viruses. (**a**) Determination of the pH threshold triggering HA-mediated membrane fusion. Vero cells were infected with VSVΔG(HA:NA:GFP) encoding the HA and NA glycoproteins of HPAIV A/Pelican/Bern/1/2022 (H5N1) along with green fluorescent protein (GFP). At 18 hours post infection, the cells were exposed for 10 minutes at 37 °C to buffers adjusted to the indicated pH values and subsequently incubated for 2 hours at 37°C in MEM medium. The cells were fixed and nuclei stained with DAPI. The cells were visualized by fluorescence microscopy (nuclei: blue fluorescence; GFP: green fluorescence). Bar size = 50 μm. Arrows point to syncytia formed. (**b-d**) Analysis of pH-dependent inactivation of H5 avian influenza viruses. HPAIV A/cattle/Texas/063224-24-1/2024 (H5N1) (**b**), LPAIV A/duck/Hokkaido/Vac-1/2004 (H5N1) (b), and LPAIV A/duck/Potsdam/1402/1986 (H5N2) (**c**) were incubated for 30 minutes at 21 °C with buffers adjusted to the indicated pH values. The buffers were subsequently replaced by MEM medium using Sephadex G25 size exclusion chromatography, and infectious virus titers determined on MDCK cells. Mean values and standard deviations of three inactivation experiments are shown. Significantly different titers compared to the treatment at pH 7.0 were calculated using the one-way ANOVA test with SIDAK’s multiple comparisons (\*\**p* < 0.01, \*\*\*\**p* < 0.0001).

### 3.4. Inactivation of avian influenza viruses in a pH-dependent manner

To investigate the impact of proton concentrations on the stability of bovine H5N1 HPAIV, the virus was incubated for 30 minutes at 21°C with buffers adjusted to different pH values. The buffer was subsequently exchanged against cell culture medium using Sephadex-G25 gel filtration columns, and infectious virus titers determined on MDCK cells by end point dilution. Using the infectious titer at pH 7.0 as a reference, we found that the infectivity of the virus was not affected when the virus was exposed to pH values in the basic range (pH 8-10) or when the pH was slightly acidic (pH 6.0) (**Fig. 3b**). A gradual reduction of the infectious titer, however, was observed when the pH was lowered stepwise from pH 6.0 to 4.0, with no infectivity left at pH 4.0. Similarly, LPAIV H5N1 was stable between the pH range 6 -10, showed a significant drop of infectious titer at pH 5.0, and complete loss of infectivity when incubated at pH 4.0 (**Fig. 3c**). However, when the experiment was performed using an H5N2 LPAIV, some residual infectivity was still observed at pH 4.0 (**Fig. 3d**), suggesting that some avian influenza virus strains might be more resistant to low pH-mediated inactivation.

### 3.5. Inactivation of H5N1 LPAIV by incubation with raw milk cheese and yoghurt

To investigate the stability of H5N1 LPAIV in fermented milk products, we produced laboratory-scale soft and semi-hard mini-cheese using raw milk, cut the cheese into small pieces and incubated them for 24 hours at 4°C with a suspension of LPAIV A/duck/Hokkaido/Vac-1/2004 (H5N1) in water. In parallel, H5N1 LPAIV was incubated for 24 hours at 4°C with buffer adjusted to either pH 7.0, pH 6.0, pH 5.0 or pH 4.0. Afterwards, the buffer was replaced by MEM medium by size exclusion chromatography, and the infectious virus titer determined on MDCK cells. When the virus was incubated with a buffer adjusted to pH 6.0, the virus titer remained at the same level as when it was incubated with a buffer adjusted to pH 7.0 (**Fig. 4a**). Incubation of the virus for 24 hours with a buffer adjusted to 5.0 resulted in a drop of infectivity by 3.3 log_10_, while incubation at pH 4.0 almost completely inactivated the virus. When H5N1 LPAIV was incubated with soft cheese with a pH of 5.0 or with semi-hard cheese (pH 5.3), infectious virus titer was reduced by 5.1 log and 3.9 log_10_, respectively. Interestingly, the incubation with both kinds of cheese resulted in a more pronounced reduction of virus titer compared to incubation with buffer of pH 5.0, indicating that factors other than pH might have contributed to this dramatic loss in infectivity. To prove whether incubation with cheese or buffers of different pH would have resulted in degradation of viral RNA, total RNA was extracted from the samples and analyzed for the presence of viral RNA genome segment 7 by RT-qPCR. Compared to virus incubated at pH 7.0, we observed a reduction of the number of RNA genome equivalents when H5N1 LPAIV was incubated at acidic conditions or with one of the cheese types (**Fig. 4b**), suggesting that virus integrity was affected and some viral RNA degraded.

**Fig. 4.**
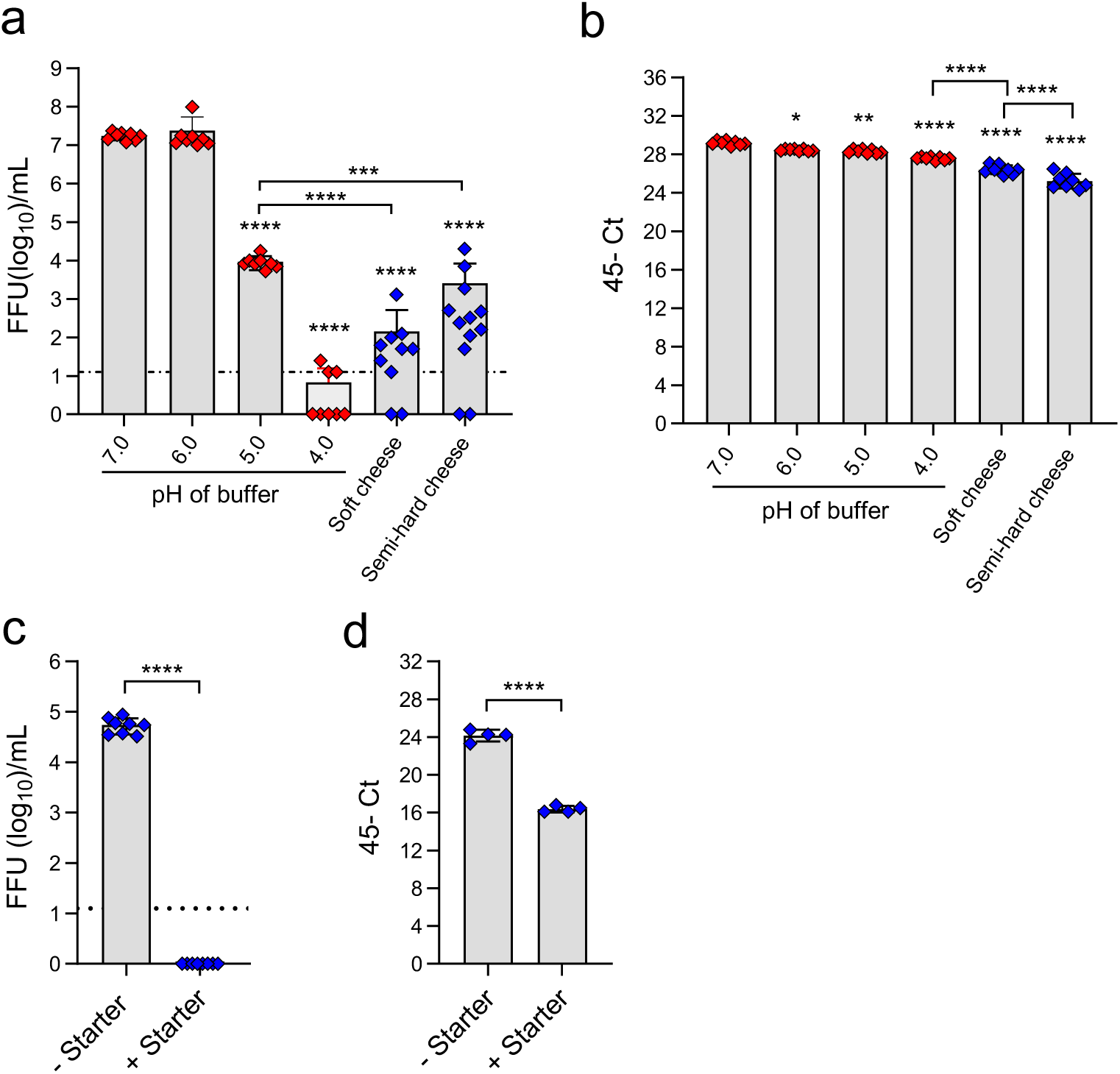
Effect of lactic acid fermentation on H5N1 LPAIV infectivity. (**a, b**) LPAIV A/duck/Hokkaido/Vac-1/2004 (H5N1) was incubated for 24 hours at 4 °C either with buffers of the indicated pH or with soft cheese or semi-hard cheese. The incubation medium was replaced afterwards by MEM medium using size exclusion chromatography. (**a**) Infectious virus titers were determined on MDCK cells. Mean values and standard deviations of 8 experiments for each condition are shown. (**b**) Detection of viral RNA by RT-qPCR. The maximum amplification cycles minus the cycle threshold number (45-Ct) are plotted. Mean values and standard deviations are shown. Significant differences were assessed by the one-way ANOVA test with Tukey’s or Dunnett’s multiple comparisons (\**p* < 0.05; \*\**p* < 0.01; \*\*\**p* < 0.001, \*\*\*\**p* < 0.0001). (**c**) Inactivation of H5N1 LPAIV during the production of yoghurt. Fresh raw milk was spiked with LPAIV A/duck/Hokkaido/Vac-1/2004 (H5N1) and incubated with or without starter culture at 42 °C. When the pH of the yoghurt was in the range of 4.2 to 4.6, and milk became thick, the yoghurt was stored overnight at 4 °C. The yoghurt was diluted with MEM medium, sterile-filtrated, and infectious virus titers determined on MDCK cells. Mean infectious titers and standard deviations for eight yoghurt samples are shown. (**d**) Detection of H5N1 RNA by RT-qPCR. Mean values and standard deviation of four samples analyzed are shown. Significant differences in virus titer were assessed using the unpaired two-tailed student t test (\*\*\*\**p* < 0.0001).

Finally, we produced yoghurt by incubating H5N1 LPAIV-spiked raw milk with or without starter culture for about 5 hours at 42°C. When the pH of the milk with starter culture was in the range of 4.2 to 4.6, the fermentation process was stopped by incubating the yoghurt overnight at 4°C. Milk to which no starter culture had been added showed a pH of 6.5. Finally, the both preparations were mixed with MEM medium, sterile-filtrated, and infectious virus titer determined by titration on MDCK cells. Using this approach, we detected significant titers of infectious virus in the yoghurt preparation without starter culture, whereas no infectious virus was found in the yoghurt preparation to which starter culture had been added and fermentation took place (**Fig. 4c**). RT-qPCR analysis of the samples revealed a significant drop in intact viral RNA in yoghurt where fermentation took place and pH dropped to pH 4.3 (**Fig. 4d**), suggesting that some viral RNA had been degraded.

## 4. Discussion

Consumption of raw milk is known to have positive effects on human health by supporting the development of a versatile gut microbiome and protecting from allergies (Bachmann et al., 2020; Kok and Hutkins, 2018; Montel et al., 2014). Nonetheless, people consuming raw milk or raw milk products from cows, goats or sheep may be exposed to bacterial pathogens such as *Campylobacter*, *Listeria*, *Brucella*, Shiga toxin-producing *Escherichia coli* and *Salmonella*, and viral pathogens such as tick-borne encephalitis virus (TBE) and hepatitis E virus (Gonzalez et al., 2022; Holzhauer and Wennink, 2023; Huang et al., 2016; Idland et al., 2022; Williams et al., 2023; Zahmanova et al., 2024). Since H5N1 highly pathogenic avian influenza virus (HPAIV) has recently been detected in the milk of lactating cows in the United States of America, this virus now needs to be added to this list of pathogens of concern. Pasteurization of milk has been shown to inactivate most of these pathogens if performed properly (Alkie et al., 2025; Cui et al., 2024; Oliver et al., 2005; Schafers et al., 2025; Spackman, Anderson, et al., 2024). Nevertheless, raw milk is still widely consumed. In addition, several types of cheese are made from raw milk. Therefore, the aim of this study was to investigate whether cheese made from raw milk containing infectious H5N1 avian influenza viruses would pose a food safety risk. As lactic acid fermentation is a key process in cheese production, we investigated the impact of two important parameters in this process, temperature and pH for viral infectivity.

First, we investigated whether temperature along with prolonged storage time would affect virus infectivity. Incubation of HPAIV and LPAIV H5N1 with milk showed that the infectious titer of these viruses remained stable for at least four weeks. Incubation with milk at 21°C resulted in a moderate decrease in infectivity over time. This decrease was more pronounced when the virus was incubated at 37°C. We also observed strain-specific differences, as LPAIV H5N1 was more stable than HPAIV H5N1 during long-term storage at these temperatures. The mechanism for these differences in temperature-dependent slow virus inactivation is not known. Inactivation of hepatitis C virus by human breast milk has been shown to be due to modification of the viral envelope by specific fatty acids in human breast milk (Pfaender et al., 2013). Presently, it cannot be excluded that a similar mechanism contributes to the slow inactivation of H5N1 avian influenza viruses when incubated with bovine milk at 21°C or 37°C.

Short-term (30 minutes) incubation of H5N1 LPAIV at elevated temperatures showed that the virus is relatively stable at temperatures up to 45 °C. The resistance of H5N1 avian influenza viruses to inactivation at this temperature may be a consequence of their adaptation to the body temperature of avian hosts, which is usually about 41.0 °C during the active phase and up to 43.8 °C when birds are highly active (Prinzinger et al., 1991). Incubation of H5N1 LPAIV viruses at 50 °C resulted in a more pronounced decrease in infectious virus titers, most likely due to temperature-mediated denaturation of viral proteins. The higher the temperature, the shorter the time required for complete virus inactivation. Pasteurization at 72 °C for 15 seconds efficiently reduced viral infectivity, consistent with previous reports (Alkie et al., 2025; Spackman, Anderson, et al., 2024). It should be noted that temperatures during raw milk cheese production are typically below 55°C, except for a few hard and extra-hard cheeses. Therefore, the temperature during lactic acid fermentation may not result in sufficient inactivation of H5N1 HPAIV in bovine milk.

Exposure of H5N1 HPAIV or LPAIV to buffers of different pH showed that the viruses were stable in the alkaline range (up to pH 10.0), but were inactivated in the acidic range, especially when the pH dropped below pH 5.0. One explanation for this acid lability is probably the role of protons in the life cycle of influenza A viruses. The dominant glycoprotein in the viral envelope, hemagglutinin (HA), has pH-dependent membrane fusion activity. This membrane protein is synthesized in the infected cell as a precursor protein that is cleaved into two subunits by cellular proteases (Bottcher-Friebertshauser et al., 2014). This post-translational modification causes HA to adopt a metastable conformation that confers fusion competence. Membrane fusion usually occurs only after the virus has been taken up by the host cell via receptor-mediated endocytosis. Acidification of the endosomes below a certain pH threshold leads then to a conformational change of the HA, which in turn initiates membrane fusion (Russell, 2014; Russell et al., 2018). In addition, protons enter the virus through the M2 ion channel in the viral envelope, leading to disruption of the M1 matrix protein layer and other protein-protein interactions (Chlanda and Zimmerberg, 2016). The pH-induced conformational change of HA is thought to be irreversible (Benhaim et al., 2020; Benton et al., 2020). Therefore, exposure of a virus particle to acidic medium prior to cell entry would result in premature denaturation of the HA protein and loss of infectivity. Following the low pH-triggered conformational change, HA is also known to be more susceptible to proteolytic attack and degradation (Puri et al., 1990), which may additionally contribute to loss in virus infectivity.

Interestingly, the pH threshold triggering HA conformational change and HA-mediated membrane fusion was found to occur at mildly acidic conditions (pH 6.0 - 6.1). This is consistent with recent data showing that membrane fusion by various isolates of H5N1 HPAIV clade 2.3.4.4b is triggered by a pH of 5.9 (Yang et al., 2025), a typical pH threshold value common to most avian influenza viruses (Russell et al., 2018). Nevertheless, complete inactivation of H5N1 HPAIV was only observed when the virus was incubated with buffer adjusted to a pH of 4.0. When the virus was incubated at a pH of 5.0, a considerable amount of infectious virus was retained (see **Fig. 3b, c, d**). This finding is consistent with the notion that membrane fusion by influenza virus may involve conformational changes of only a small HA fraction (Remeta et al., 2002). Another fraction of HA may not completely unfold but may stay at transition stages which are not irreversible and allow HA to fold back into the metastable conformation when the pH is raised again. The incubation of H5N1 LPAIV with laboratory-scale soft cheese (pH 5.0) or semi-hard mini-cheese (pH 5.3), respectively, had a stronger inactivating effect on H5N1 LPAIV infectivity compared to the incubation with buffer adjusted to pH 5.0. The reason for this phenomenon is not yet understood. However, it is known that lactic bacteria produce proteolytic enzymes that contribute to cheese maturation (Kok, 1991; Law and Kolstad, 1983). On the other hand, it is well established that the low pH-induced conformational change of HA renders the molecule more susceptible to proteolytic degradation (Puri et al., 1990). We therefore hypothesize that H5N1 HPAIV remains infectious in raw milk even after the pH has dropped to pH 5.0 will further lose infectivity due to bacterial proteases involved in cheese maturation. Depending on the recipe of cheese-making, this process can last from 2 weeks for some types of soft cheeses up to 2 years for certain extra-hard cheeses, and usually takes place at temperatures between 10-16 °C. More detailed experiments with longer ripening times will show how fast this inactivation process is and whether there are differences in the inactivation kinetics, depending on the type of cheese and used recipe.

Based on the data presented in this study, a preliminary risk assessment of the consumption of fermented raw milk products is possible. We found that the temperatures which are used for the manufacture of most raw milk cheeses, with exception of some hard and extra-hard cheeses, are not sufficient to completely inactivate H5N1 viruses. In addition, the temperatures at which cheese ripening occurs for extended periods of time will likely not affect infectious titers to a significant degree. However, the decrease in pH during lactic acid fermentation leads to a significant reduction in infectious titer. Bacterial proteases involved in cheese maturation may further reduce the infectivity of avian influenza viruses in raw milk products. We therefore conclude that the proton concentration is the most important parameter for the inactivation of avian influenza viruses in raw milk dairy products. Depending on the recipe, acidification below a pH of 5.3 leads to a large reduction of viral titers, while acidification below pH of 4.3 fully inactivates these viruses.

## CRediT authorship contribution statement

**Nicole Lenz-Ajuh**: Conceptualization, Investigation, Writing – review and editing; **Leonie Rau**: Data curation, Formal analysis, Investigation, Visualization; **Lisa Butticaz**: Data curation, Formal analysis, Investigation; **Bettina Zimmer**: Data curation, Investigation, Methodology; **Vincent Beuret**: Methodology, Resources; **Florian Loosli**: Methodology, Resources; **Jan-Erik Ingenhoff**: Conceptualization; **Barbara Wieland**: Conceptualization, Funding acquisition, Writing – review and editing; **Gert Zimmer**: Conceptualization, Data curation, Formal analysis, Funding acquisition, Investigation, Methodology, Project administration, Supervision, Validation, Visualization, Writing – original draft, Writing – review and editing.

## Acknowlegements

We are grateful to the Federal Food Safety and Veterinary Office (FSVO) and the Swiss Expert Committee for Biosafety (SECB) for financial support. We like to thank Yoshihiro Sakoda and Timm Harder for their generous gifts of low-pathogenic avian influenza viruses.

## Data availability

All data pertaining to this study are presented in the paper and its figures. Source data are available on request.

